# TRPC5-Ca_V_3 complex mediates Leptin-induced excitability in hypothalamic neurons

**DOI:** 10.1101/2020.07.21.214296

**Authors:** Paula P. Perissinotti, Elizabeth Martínez-Hernández, Erika S. Piedras-Rentería

**Author notes:** Authors contributed equally to this work (Paula P. Perissinotti, Elizabeth Martínez-Hernández). Corresponding author: Erika S. Piedras-Rentería, Cell and Molecular Physiology Department, Loyola University Chicago, Stritch School of Medicine, 2160 S. First Ave., Maywood IL 60153., **Email:**. Instituto de Fisiología, Biología Molecular y Neurociencias (IFIBYNE, UBA-CONICET), Ciudad Universitaria, Buenos Aires, C1428EHA, Argentina.

## Abstract

Leptin regulates hypothalamic POMC^+^ (pro-opiomelanocortin) neurons by inducing TRPC (Transient Receptor Potential Cation) channel-mediate membrane depolarization. Here we assessed the role of T-type channels on POMC neuron excitability and leptin-induced depolarization *in vitro*. We demonstrate T-type currents are indispensable for both processes, as treatment with NNC-55-0396 prevented the membrane depolarization and rheobase changes induced by leptin in cultured mouse POMC neurons. Furthermore, we demonstrate TRPC1/C5 channels and Ca_V_3.1 and Ca_V_3.2 channels co-exist in complex. The functional relevance of this complex was corroborated using intracellular Ca^2+^ chelators; intracellular BAPTA (but not EGTA) application was sufficient to preclude POMC neuron excitability by preventing leptin-induced calcium influx through TRPC channels and T-type channel function.

We conclude T-type channels are integral in POMC neuron excitability. Leptin activation of TRPC channels existing in a macromolecular complex with T-type channels recruits the latter by locally-induced membrane depolarization, further depolarizing POMC neurons, triggering action potentials and excitability.

## INTRODUCTION

Leptin regulates energy homeostasis and serves as a satiety afferent signal in the homeostatic control of adipose tissue mass (Harvey & Ashford, 2003; Schwartz, Woods, Porte, Seeley, & Baskin, 2000), reducing food intake, increasing energy expenditure and regulating the reward value of nutrient (Ahima & Flier, 2000; Domingos et al., 2011; Williams & Elmquist, 2012). Leptin exerts its physiological action through its specific receptor (LRb), which is highly expressed in hypothalamus and other brain areas (Shioda et al., 1998). Leptin’s effects on hypothalamic homeostatic feeding circuits are well established (Kenny, 2011); it negatively modulates orexigenic agouti-related peptide (AgRP)/neuropeptide Y (NPY) neurons by Kv2.1 channel-mediated membrane hyperpolarization (Baver et al., 2014). In contrast, anorexigenic POMC positive neurons are depolarized by leptin (Cowley et al., 2001). This depolarization is mediated via a Jak2-PI3 kinase-PLCγ pathway that ultimately activates TRPC channel activity (Qiu et al., 2010). Similarly, a subset of POMC neurons in the arcuate nucleus responsive to serotonin *via* the 5-HT_2C_ receptor are also activated *via* TRPC channels, suggesting TRPC channels are a common signaling mechanism mediating anorexigenic signaling in the hypothalamus (Sohn et al., 2011). TRPC5 channels are also the molecular mediators of the acute leptin and serotonin effect in POMC neurons (Gao et al., 2017).

The role of TRPC channels in POMC neuron excitability is clearly established; however, it is not known whether their activity alone is sufficient to trigger excitability. Here we used cultured hypothalamic neurons from mice to characterize the role of T-type Ca^2+^ channels and leptin-induced POMC neuron excitability. Our data demonstrate T-type channels are necessary for POMC neuron excitability, by being involved in the excitatory cascade induced by leptin in these neurons. Blockade of either TRPC or T-type channel function prevents the effect of leptin on hypothalamic neuron excitability. Moreover, we demonstrate that: a) TRPC1 and TRPC5 channels co-immunoprecipitate with T-type channels in the hypothalamus, b) TRPC-T-type channel complexes exist in a functional microdomain, and c) TRPC-induced depolarization in these domains triggers neuronal excitability *via* recruitment of T-type channels. Thus, this study confirms T-type channels constitute a target to modulate leptin-activated neurons and their functions, such as energy balance and food intake.

## RESULTS

### T-type calcium currents are necessary for the excitability of leptin-responsive hypothalamic neuron

Hypothalamic cultures were studied from 8-10 days in vitro (DIV). Immunocytochemistry analysis (IC) showed that POMC and NPY neurons were present in the culture, with POMC neurons (POMCN) being the majority (85 *vs*. 15%, n= 182; *P* < 0.05, Z-test) (Fig. 1A and B). Electrophysiological experiments confirmed 80% of all neurons tested were Leptin (Lep)-activated (Fig. 1C and D), consistent with the IC data. The properties of these neurons were consistent with POMC neuron responses: their resting potential was slightly depolarized, their rheobase was decreased and the number of action potentials (APs) was increased upon application of 100 nM Lep (Table 1). In contrast, Lep treatment increased the rheobase and decreased AP number in 5% of cells, consistent with an NPY phenotype. 15 % of neurons tested where neither responsive nor inhibited by leptin. We also confirmed the presence of Ca_V_3.1 and Ca_V_3.2 channels in POMC^+^ neurons by immunocytochemistry (Figure 2A).

**Figure 1.**
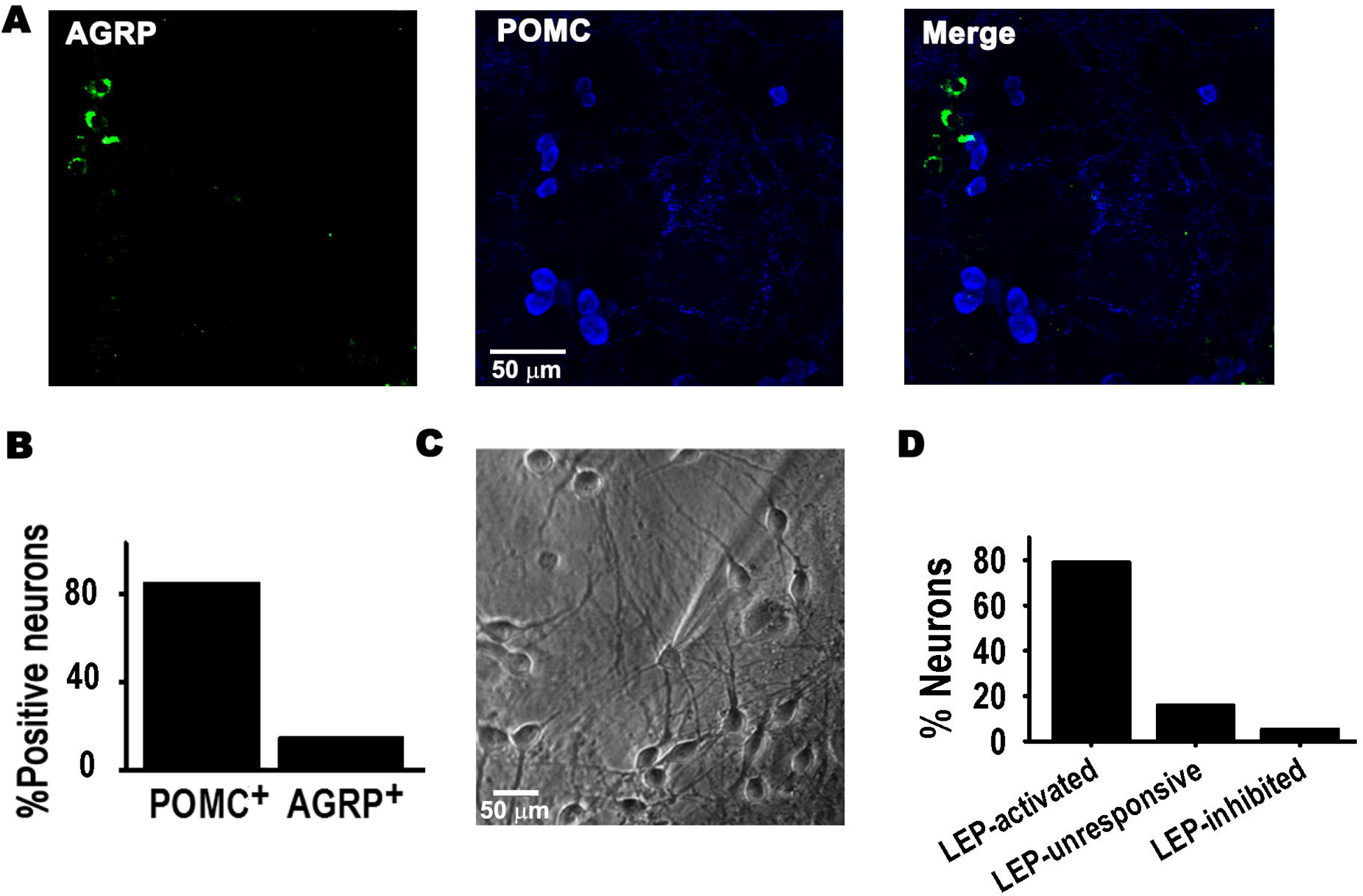
POMC neurons activated by leptin in culture. (A) Example of confocal images showing neurons positive for AGRP (green) or POMC (blue) antibodies in hypothalamus culture at 8-10 DIV. **(B)** Percentage of neurons positive for AGRP or POMC antibodies (n= 21) **(C)** Microscopic image of cultured hypothalamic neuron at 8 DIV. **(D)** Percentage of WT neurons activated or inhibited by 100 nM leptin (n= 19), using a 20 pA/s depolarizing ramp stimulus.

**Figure 2.**
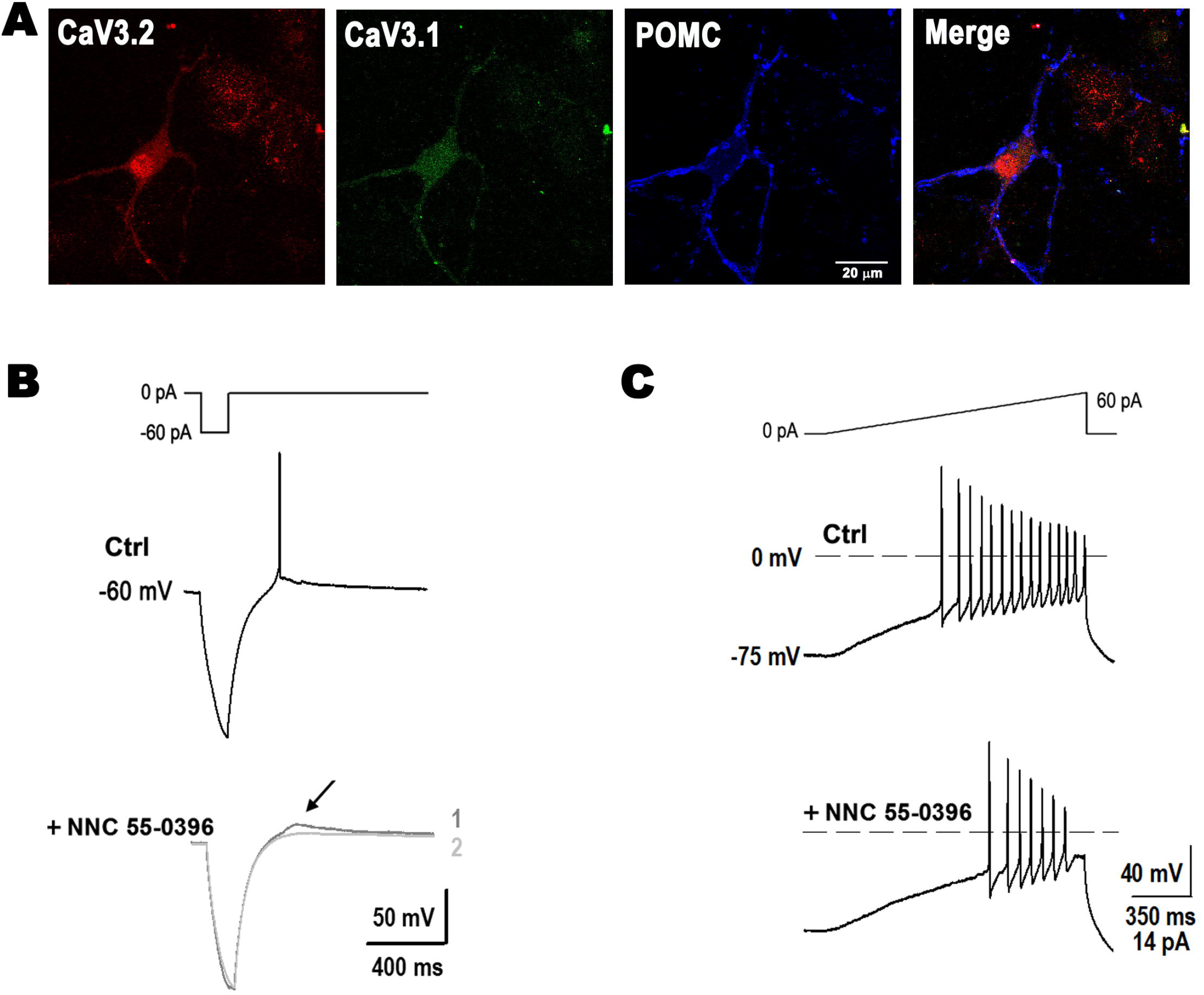
T-type calcium currents mediate membrane excitability in POMC neurons. **(A)** Example of confocal images showing the expression of T-type α_1H_ (Ca_V_3.2, red) and α_1G_ (Ca_V_3.1, green) channels in a POMC neuron (blue). Bar size, 20 μm. **(B)** Post-inhibitory rebound before (Ctrl) and after perfusion with 10 μM NNC 55-0396. Arrow, DTD (#1) disappeared after longer perfusion with NNC 55-0396 (2). **(C)** Application of NNC 55-0396 reduced membrane excitability (V_H_= −75 mV, 40 pA/s ramp).

**Table 1.**
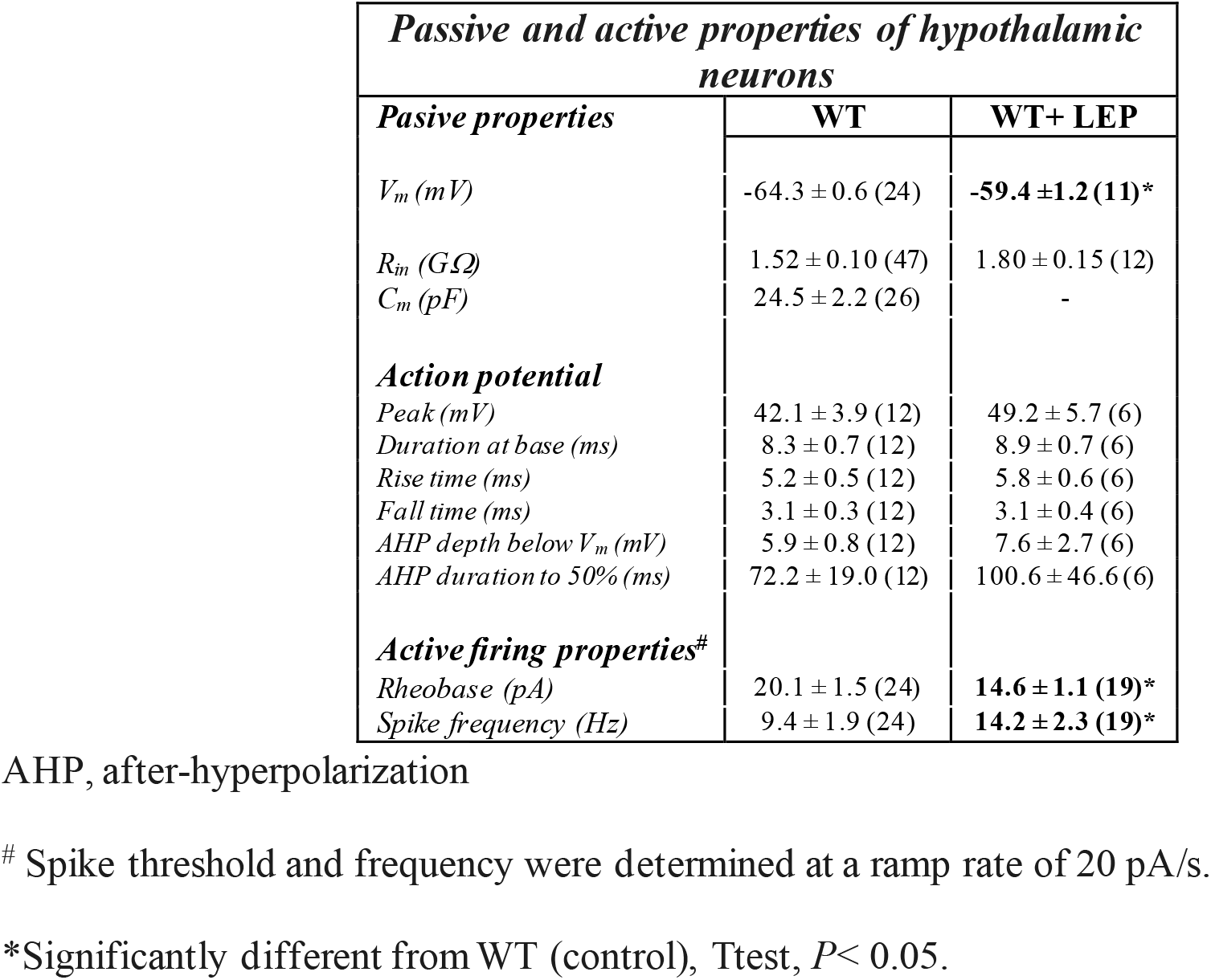
Passive and active properties of hypothalamic neurons

| *Passive and active properties of hypothalamic neurons* |  |  |
| --- | --- | --- |
| *Passive properties* | **WT** | **WT+ LEP** |
| $V_m$ (mV) | $-64.3 \pm 0.6$ (24) | **$-59.4 \pm 1.2$ (11)\*** |
| $R_{in}$ (GΩ) | $1.52 \pm 0.10$ (47) | $1.80 \pm 0.15$ (12) |
| $C_m$ (pF) | $24.5 \pm 2.2$ (26) | - |
| *Action potential* |  |  |
| Peak (mV) | $42.1 \pm 3.9$ (12) | $49.2 \pm 5.7$ (6) |
| Duration at base (ms) | $8.3 \pm 0.7$ (12) | $8.9 \pm 0.7$ (6) |
| Rise time (ms) | $5.2 \pm 0.5$ (12) | $5.8 \pm 0.6$ (6) |
| Fall time (ms) | $3.1 \pm 0.3$ (12) | $3.1 \pm 0.4$ (6) |
| AHP depth below $V_m$ (mV) | $5.9 \pm 0.8$ (12) | $7.6 \pm 2.7$ (6) |
| AHP duration to 50% (ms) | $72.2 \pm 19.0$ (12) | $100.6 \pm 46.6$ (6) |
| *Active firing properties<sup>#</sup>* |  |  |
| Rheobase (pA) | $20.1 \pm 1.5$ (24) | **$14.6 \pm 1.1$ (19)\*** |
| Spike frequency (Hz) | $9.4 \pm 1.9$ (24) | **$14.2 \pm 2.3$ (19)\*** |

AHP, after-hyperpolarization
<sup>#</sup> Spike threshold and frequency were determined at a ramp rate of 20 pA/s.
\*Significantly different from WT (control), Ttest, $P < 0.05$ .

We first studied the role of T-type currents on POMC neuron excitability by post-inhibitory rebound experiments (PIR). Transition from hyperpolarized to depolarized membrane potentials resulted in T-type channel activation and AP firing in control conditions, which was completely prevented by application of 10 μM of T-type channel blocker NNC-55-0396 (n= 5) (Fig. 2B). A delayed transient depolarization (DTD) due to calcium entry can be seen in the trace immediately after the application of the blocker (bottom, trace 1, arrow), which quickly disappeared upon continued perfusion with the drug (trace 2). Membrane excitability was also quantified by assessing the rheobase and the number of APs with a current ramp stimulus from V_H_= −75 mV to activate T-type calcium channels (ramp rates of 20 and 40 pA/s). Neuronal membrane excitability elicited with 20 pA/s ramp rates (20.1 ± 1.5 pA, 7.8 ± 1.6 APs, n= 24, 20 pA/s) was not significantly altered by the addition of NNC-55-0396 (23.8 ± 2.3 pA, 6.2 ± 1.8 APs, n= 5, *P*> 0.05). However, application of NNC 55-0396 after stimulation with ramp rates of 40 pA/s elicited reduced excitability (increased rheobase) (36.9 ± 3.3 pA) and decreased number of APs to 9.6 ± 3.7 (n= 5) compared to control (rheobase 24.9 ± 2.3 pA, 18.6 ± 1.9 APs (n= 24), *P*< 0.05, Fig. 2C). These results corroborate the role of T-type channels in membrane excitability.

### T-type (LVA) current assessment

Analysis of Ca^2+^ current properties revealed three distinct neuron groups, as seen in the I-V curves depicted in Fig. 3A. Of all neurons sampled (n= 28), 14.3% expressed only high-voltage activated (HVA) currents; 46.4% of neurons expressed both HVA and LVA currents at low levels (LD, I_LVA peak_〈6-pA/pF), and 39.3% expressed low HVA and high LVA current density levels (HD, I_LVA peak_ from 6 to 20 -pA/pF) (Panel 3A-B).

**Figure 3.**
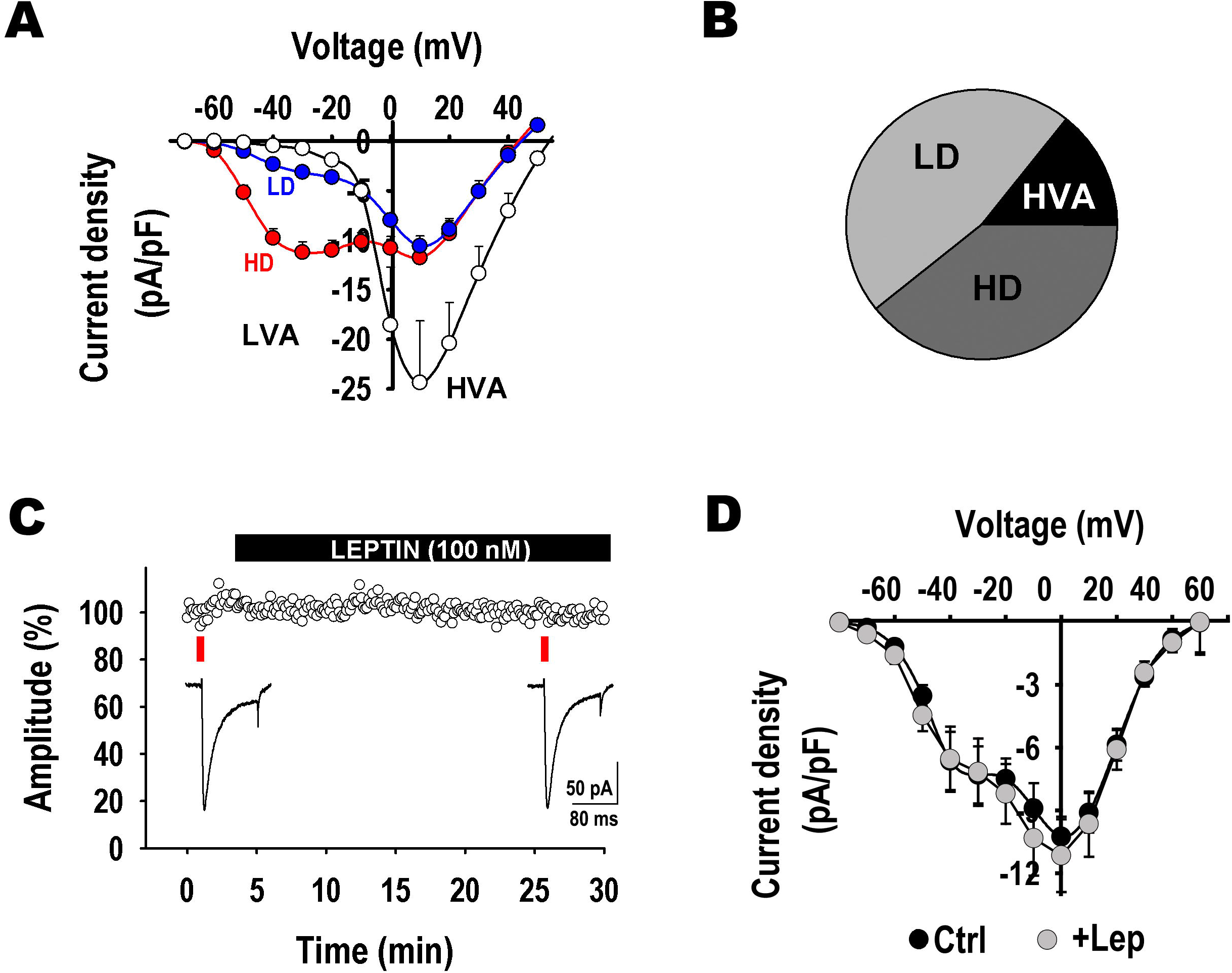
Leptin has no direct effect on T-type current density. **(A)** I-V curves depict three distinct neuron groups (n= 28): empty circles, neurons expressing only high-voltage activated (HVA) currents; blue circles, neurons expressing low HVA and low LVA current density levels (LD, I_LVA peak_〈6 -pA/pF); and red circles, neurons expressing low HVA and high LVA current density (HD, I_LVA peak_ from 6 to 20 -pA/pF). **(B)** Pie chart denotes the percentual number of neurons in each group described in A. **(C)** Acute incubation with Lep (100 nM) does not alter LVA current density elicited by a step voltage from −90 to −30 mV (n= 3). **(D)** I-V curves before and after the application of 100 nM leptin in the recording chamber (n= 8, *P*>0.05 at - V_T_=30 mV).

Given that leptin is known to exert direct effects on voltage-gated calcium currents (Takekoshi et al., 2001; Wang et al., 2008), we assessed whether acute incubation with Lep altered LVA currents. As seen in Figure 3C-D, application of 100 nM leptin in the recording chamber did not alter the current density or their kinetics of activation or inactivation (n= 8, *P*> 0.05 at -V_T_= 30 mV).

### T-type calcium currents are necessary for the leptin-induced excitability response

We next assessed whether T-type channels are part of the cellular pathway of neurons depolarized by leptin. Treatment of POMC neurons with Lep induces enhanced PI3K activity and ultimately results in activation of transient receptor potential cation channels (TRPC) and a subsequent discrete resting membrane depolarization (Qiu, Fang, Bosch, Ronnekleiv, & Kelly, 2011; Qiu et al., 2010; Qiu et al., 2014). The presence of TRPC channels in POMC^+^ neurons was confirmed by immunocytochemistry (Figure 2A). We then assessed the effect of Lep on excitability using depolarizing current ramps. Figure 4B shows an example of neuronal firing evoked in a POMC neuron at ramp rates of 20 pA/s. As anticipated, Lep treatment on its own decreased the rheobase from 18.4 to 14.1 pA and increased the number of action potentials from 6 to 11. The summary of effects of Lep on neurons is depicted in Fig. 4C and also in Table I. Similar to the application of NNC-55-0396, addition of 2-APB alone did not affect neuronal membrane excitability *per se* (20.1 ± 0.4 pA, 7.7 ± 3.1 APs, n= 4, *P*> 0.05). However, the effect of leptin was abolished in the presence of 100 μM of TRPC channel blocker 2-APB, resulting in 24.5 ± 4.3% increase in rheobase and 48.0 ± 5.3% decreased number of spikes, compared to the leptin-treated group; corroborating the role of these channels in the leptin-signaling cascade (Fig. 4C). Interestingly, the effect of Lep was also prevented solely by the application of the T-type channel blocker NNC-55-0396 (*P*< 0.05, ANOVA, Fig. 4C), demonstrating T-type channel recrutiment is necessary in this pathway, likely downstream of TRPC channel activity, as summarized in the cartoon in Fig. 4D.

**Figure 4.**
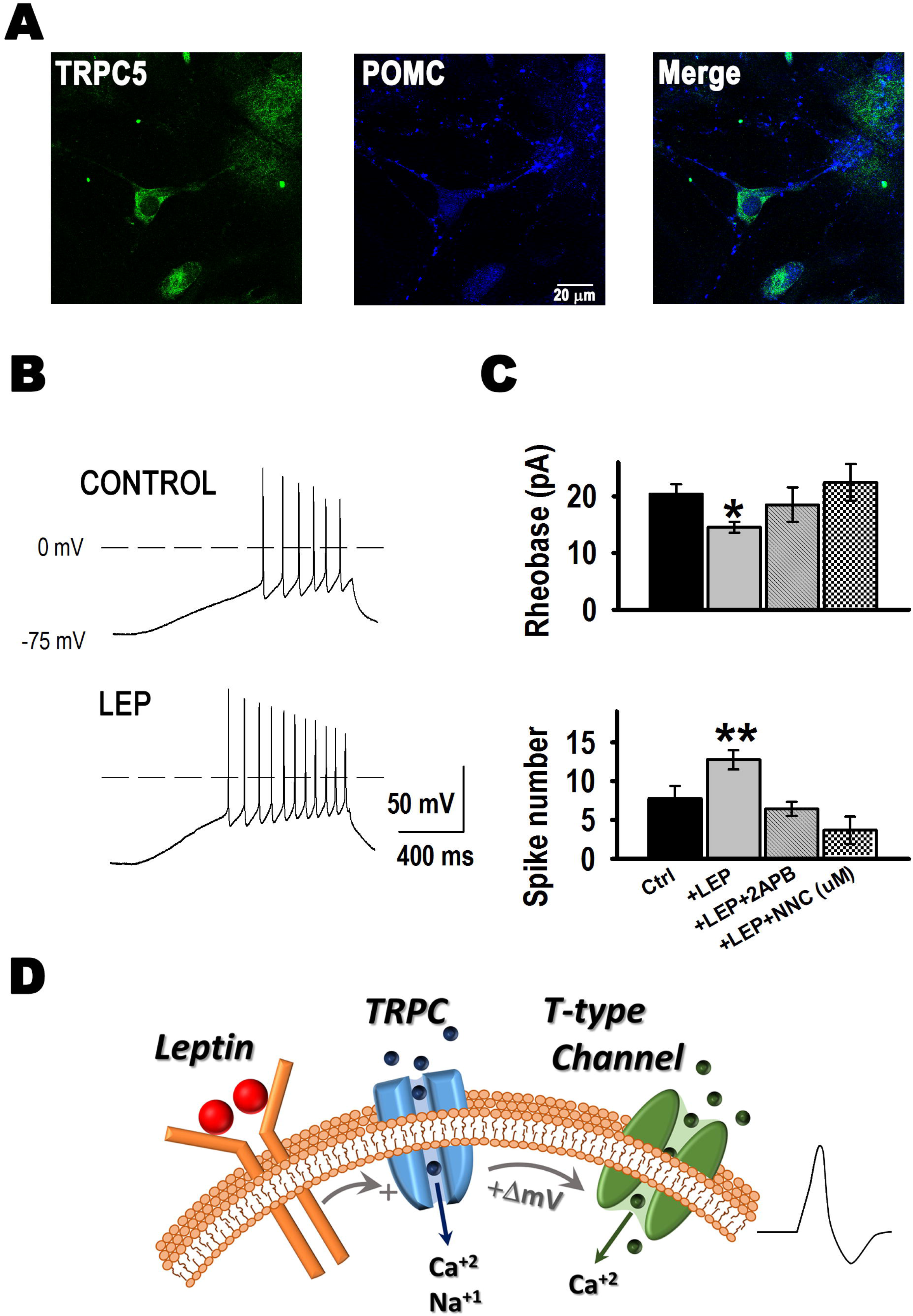
Leptin increases excitability in cultured hypothalamic neurons. **(A)** Example of confocal images showing the expression of TRPC5 channels (green) in POMC neurons (blue). **(B)** Examples of neuronal firing in control and Lep-treated neurons elicited from VH= −75 mV by a 20 pA/s ramp. **(C)** Quantification of rheobase (pA) and number of action potentials in control neurons (n= 26), neurons treated with Lep (100 nM, n= 19), neurons treated with Lep and TRPC channel blocker 2-APB (100 μM, n= 5), and neurons treated with Lep and T-type channel blocker NNC 55-0396 (10 μM, n= 5). *Significantly different from control and +Lep+NCC, **significantly different from all others, *P*< 0.05, ANOVA. **(D)** Updated model of leptin’s cellular signaling. Lep binding to its receptor results in activation of the PI3 kinase signaling pathway and subsequent activation of TRPC channel. TRPC channel activity induces a small membrane depolarization that recruits T-type channel activity, increasing hypothalamic neuron excitability.

### T-type calcium currents exist in a functional microdomain complex with TRPC1/5 channels

Calcium channels are known to co-exist with other channels in functional complexes (Robitaille, Garcia, Kaczorowski, & Chariton, 1993; Vivas, Moreno, Santana, & Hille, 2017), notably T-type channels form protein complexes with members of the potassium channel family such as Kv4, KCa3.1, and KCa1.1 (Anderson et al., 2010; Rehak et al., 2013), which ensures rapid potassium channel activation thanks to their proximity with Ca_V_3 channels within the microdomain. Thus, we explored whether TRPC and T-type channels exist in a complex. TRPC5 or TRPC1/5 complexes are thought to be the physiological mediators of leptin’s effects in the hypothalamus (Strübing, Krapivinsky, Krapivinsky, & Clapham, 2001), therefore we assessed whether Ca_V_3 channels co-precipitated with TRPC5 channels in mouse whole brain and hypothalamus samples. Figure 5A shows that Ca_V_3.1 channels co-precipitate with TRPC5 channels (left), and similarly, TRPC 5 channels co-precipitate with Ca_V_3.1 channels (right panel). Ca_V_3.2 channels also co-precipitate with TRPC5 channels (Figure 5B, left), and TRPC5 interact with Ca_V_3.2 (right). We also determined that TRPC1 is detected in samples co-precipitated with TRPC5 and Ca_V_3.1 (left) or Ca_V_3.2 (right) (Fig. 5C), confirming a TRPC multimer formed by TRPC1/5 is present in the hypothalamus.

**Figure 5.**
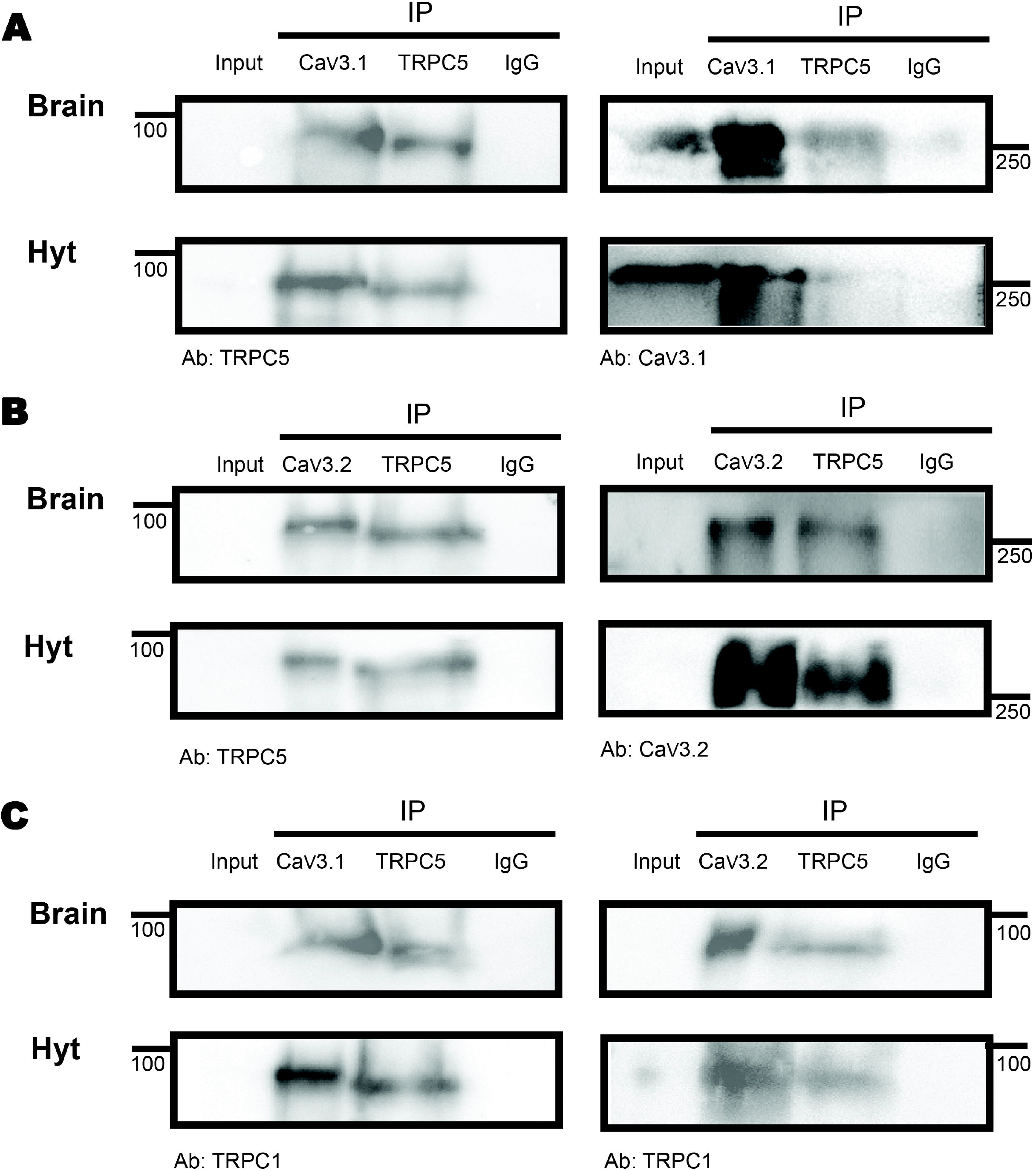
Ca_V_3 co-precipitation with TRPC5. Immunoprecipation of brain and hypothalamus (Hyt) extracts using Ca_V_3.1 **(A)** or Ca_V_3.2 **(B)** and TRPC5 in the same samples show their mutual interaction. **(C)** IP of brain and Hyt samples using Ca_V_3.1 (left) or Ca_V_3.2 (right) and TRPC5 show their interaction with TRPC1. IgG was used as IP negative control in all cases, (n=3 for all panels).

If T-type calcium channels co-exist in a complex with TRPC1/5, it is possible that membrane depolarization induced by TRPC channel-mediated cation influx could recruit T-type channel activity. As the ionic selectivity for TRPC1/5 channels is almost 1:1 for Na^+^ and Ca^2+^, Ca^2+^ ions contribute to ~66% to TRPC1/5-mediated depolarization. Accordingly, chelation of Ca^2+^ influx with a fast chelator that binds Ca^2+^ closer to the mouth’s pore (BAPTA-AM, (K_ON_ = 6 x10^8^) or with a slower chelator (EGTA-AM, K_ON_ = 1.5 x10^6^) should allow us to discern whether the TRPC1/5 and T-type channel complex function within a microdomain (Fig. 6A). We incubated cultured hypothalamic neurons with either 10 μM BAPTA-AM or EGTA-AM for 30 minutes at 37 °C prior to whole-cell current clamp to assess excitability. Figure 6B shows example traces of APs elicited with a 20 pA/s ramp protocol under control conditions (top trace), and after the addition of 100 nM Lep in the presence of BAPTA (middle) or EGTA (bottom). The presence of intracellular BAPTA or EGTA did not alter the baseline excitability response compared to control (BAPTA, n= 9; EGTA n= 4; P> 0.05, ANOVA; the rheobase and AP number values for control and +Lep have been reproduced in Fig 6B from Fig. 4 for comparison). However, addition of 100 nM Lep in the presence of BAPTA did not elicit significant changes in either AP or rheobase (n= 9, *P*> 0.05, Paired t-test), whereas it caused the expected excitability increase in the presence of EGTA (n= 4, *P*< 0.05, Paired t-test), indicating Ca^2+^ influx via TRPC1/5 and subsequent depolarization is either necessary for the TRPC1/5 complex activity (Strübing et al., 2001) or important for the recruitment of Ca_V_ channels and subsequent AP generation. The latter hypothesis is supported by the fact that Lep application and consequent TRPC1/5 channel activation induced a ~6 mV depolarization (Fig. 6C), which induces changes in T-type channel availability (SSI) and open probability (SSA) (Fig. 6D top, red line), that leads to a voltage right-shift into the T-type window current range, resulting in an increase in the steady state T-type calcium current from 30 to 60% (Figure 6D, bottom).

**Figure 6.**
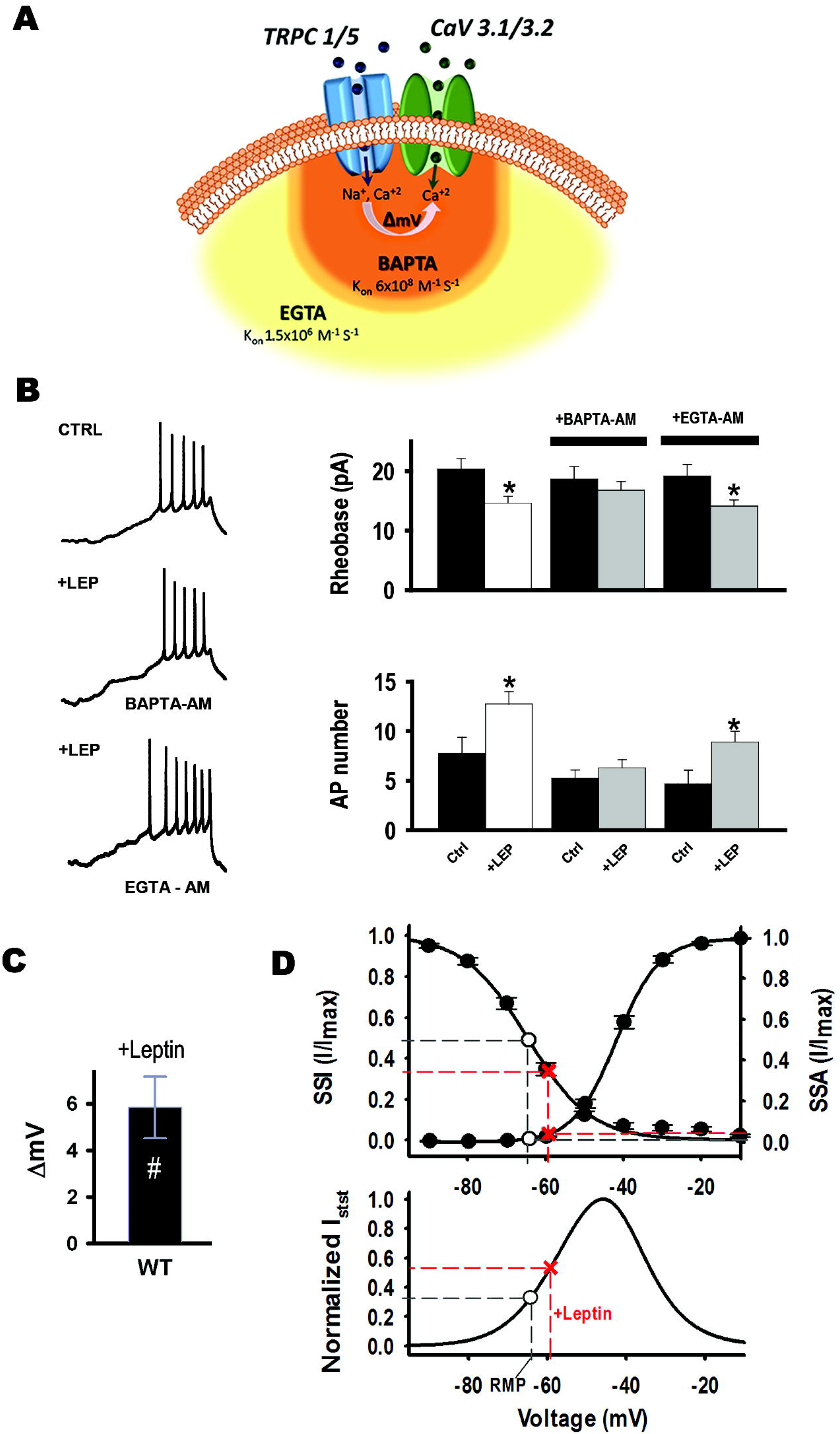
Ca^2+^ influx via TRPC1/5 depolarizes and recruits Ca_V_ channels. **(A)** Cartoon depicts that T-type calcium channels co-exist in a complex with TRPC1/5. Membrane depolarization induced by TRPC channel-mediated cation influx recruits T-type channel activity. Chelation of Ca^2+^ influx with a fast chelator that binds Ca^2+^ closer to the mouth’s pore (BAPTA-AM, (K_ON_ = 6 x10^8^) is represented in orange. Chelation of Ca^2+^ influx with a slower chelator (EGTA-AM, K_ON_ = 1.5 x10^6^) is represented in light orange. **(B)** Cultured hypothalamic neurons were incubated with either 10 μM BAPTA-AM (n= 9) or EGTA-AM (n= 4) for 30 minutes at 37 °C prior to whole-cell current clamp to assess excitability. Left: Example traces of APs elicited with a 20 pA/s ramp protocol under control conditions (top trace), and after the addition of 100 nM Lep in the presence of BAPTA (middle) or EGTA (bottom). Right: Rheobase and AP number values for each treatment. *Significantly different from control, *P*< 0.05, Paired t-test. **(C)** Lep application induced a ~6 mV depolarization (n= 5). #Significantly different from control, *P*< 0.05, Paired t-test. **(D)** T-type channel availability (SSI) and open probability (SSA) as function of voltage. Empty circles indicate the availability and the open probability of T-type channels at the resting membrane potential (RMP). Red crosses indicate these steady sate properties after Lep (100 nM) incubation. Window currents were theoretically calculated from SSA and SSI curves (see Methods). Steady state current (Normalized Isst) at the RMP (empty circle) increased after leptin incubation (red cross).

## DISCUSSION

This work demonstrates T-type currents are indispensable in POMC neuron excitability and that their function is essential, downstream of TRPC channel function, for transduction of the leptin cascade.

Several hypothalamic nuclei express T-type channels, which help control neuronal excitability. For instance, T-type currents are known to drive NPY neuron oscillatory activity in hypothalamic respiratory centers (van den Top, Lee, Whyment, Blanks, & Spanswick, 2004). GNRH-positive neurons express all three channel isoforms (α_1I_ > α_1H_ > α_1G_), and T-type currents are modulated by 17-ß estradiol in these cells (Bosch, Hou, Fang, Kelly, & Ronnekleiv, 2009; Li et al., 2009; Zhang, Renaud, & Kolaj, 2009). α_1G_ T-type channels have a confirmed role linking thalamocortical central regulation of wakefulness and body weight (Uebele et al., 2009), and they have been proposed as potential therapeutic targets for treating obesity (Chemin, Monteil, Perez-Reyes, Nargeot, & Lory, 2001; Chorvat, 2013; Uebele et al., 2009). T-type antagonists prescribed for epilepsy, depression, obsessive-compulsive disorder and bulimia nervosa can cause loss of appetite as a side effect (Cookson & Duffett, 1998; Traboulsie, Chemin, Kupfer, Nargeot, & Lory, 2006; Wilfong & Willmore, 2006). Hyperpolarization-induced removal of T-type channel inactivation allows for their stimulation by small depolarizations near the resting potential, rendering T-type currents optimal for regulating excitability under physiological conditions near resting state. We studied the role of T-type current function in POMCN excitability. We demonstrated T-type currents mediate excitability in POMC hypothalamic neurons in culture.

Although Lep has many functions, including effects on control of hormone release, immune system, vasculature, development and higher cognition (Domingos et al., 2011; Farr, Sloan, Keane, & Mantzoros, 2014; Friedman, 2004; Haynes, Morgan, Djalali, Sivitz, & Mark, 1999; Kim & Moustaid-Moussa, 2000; Myers, Cowley, & Munzberg, 2008); one of its prominent roles is as effector of the negative feedback loop, supporting homeostatic control of energy and food intake, and adipose tissue mass (Ahima et al., 1996). The majority of neurons in our hypothalamic cultures were Lep-activated, in line with a higher abundance of POMC positive neurons detected by IC. In these neurons, Lep induces electrical activity and depolarization *via* binding to LRb, activation of Janus tyrosine kinase 2 and the downstream activation of phosphatidylinositol 3 kinase (PI3K), resulting in activation of transient receptor potential cation channel activity (TRPC) (Harvey, 2007; Qiu et al., 2011; Qiu et al., 2010; Qiu et al., 2014). Indeed, Lep application to WT neurons *in vitro* also resulted in membrane depolarization and increased neuronal excitability. Treatment with the TRPC channel blocker 2-APB after Lep application decreased excitability as measure by the increased rheobase and decreased number of spikes, as expected. However, we found that in addition to TRPC channel activation, T-type channel activity is also essential in this pathway. Blockade of T-type channels with NNC 55-0396 completely prevented the effect of Lep, even when TRPC channel activity remained intact. Our data showed T-type channel function is essential in the response to Lep, downstream of TRPC channel activation; thus T-type currents are likely recruited by discrete changes in membrane depolarization induced by TRPC channels and are the final mediator of triggered activity. Furthermore, we demonstrate TRPC1/C5 channels and Ca_V_3.1 and Ca_V_3.2 channels exist in complex. TRPC channels are known to assemble in multiprotein complexes that include various key Ca^2+^ signaling proteins within Ca^2+^ signaling microdomains (Ambudkar, 2006). For instance, TRPC1, 3, 4, 5, 6, and 7 isoforms can form a macromolecular complex with the α_1C_ subunit of the L-type voltage-gated calcium channel (Ca_V_1.2) in atria and ventricle of developing heart (Sabourin, Robin, & Raddatz, 2011). Here we show that co-immunoprecipitation experiments show TRPC5 interacts with Ca_V_3.1 and Ca_V_3.2 channels. The functional activity of this complex was corroborated using intracellular calcium chelators; prevention of leptin-induced calcium influx through TRPC channels by intracellular BAPTA (but not EGTA) was sufficient to preclude POMC neuron excitability, due to 1) prevention of intracellular Ca^2+^-dependent potentiation of the TRPC channel complex (Strübing et al., 2001) and 2) by decreasing membrane depolarization in the Ca_V_3-TRPC complex microdomain by ~66%, as TRPC1/5 channel heterodimers display a permeability (P) P_Na_/P_Ca_ ~ 0.95 (Robitaille et al., 1993; Vivas et al., 2017).

Overall, our results show T-type channel activity is necessary for hypothalamic POMC neuron excitability and identify T-type channels as possible additional drug targets for leptin-mediated functions, such as metabolic energy regulation and control of food satiety.

## MATERIALS AND METHODS

The animal protocols used in this study were reviewed and approved by an independent Institutional Animal Care and Use Committee (IACUC). WT and EGFP-POMC^+^ mice were fed on an *ad libitum* standard commercial pellet diet.

### Cell Culture

Whole hypothalamus were dissected from newborn mice (P0) and cultured as described in (P. P. Perissinotti et al., 2014) and plated at a density of 25,000-35,000/coverslip and kept in a 5% CO_2_ humidified atmosphere at 37°C.

### Immunocytochemistry (IC)

WT neurons at 9-11 DIV were prepared as described (P. P. Perissinotti et al., 2014). Primary antibody dilutions were: POMC 1:200 (Novus); AgRP, 1:200 (Sta. Cruz), Ca_V_3.1 1:200 (Alomone) and 1:50 (Millipore), Ca_V_3.2 1:5 (supernatant, NeuroMab) and 1:200 (Sta. Cruz); TRPC1 1:100 & TRPC5 1:120 (Alomone). Secondary antibodies: Alexa-488, −594 and −647, 1:2,000 (Molecular Probes, Eugene, OR). Image acquisition was done using a Multiphoton Leica TCS SP5 microscope and analyzed with ImageJ freeware (NIH) (Schneider, Rasband, & Eliceiri, 2012).

### Immunoprecipitation (IP)

Crude membrane preparations were obtained using standard protocols, a fraction of the sample was reserved before immunoprecipitation (input); the remaining volume was divided up in equal parts for all experiments. Samples were then processed by addition of the primary antibody and incubation for 1 h at 4°C [antibodies: α1H (Santa Cruz Biotechnology, Inc., Santa Cruz, CA, USA), α1G (Millipore); TRPC5 (Alomone), TRPC1 (Santa Cruz Biotechnology, Inc., Santa Cruz, CA, USA), and IgG (Alpha Diagnostics, San Antonio, TX)], followed by overnight incubation with protein A/G agarose beads (Biovision, Mountain View, CA) on a shaking plate at 4°C. The samples were washed and then precipitated in 0.1 M glycine pH 3.5 and neutralized with 0.5 M Tris?Cl and 1.5 M NaCl pH 7.4 before SDS-PAGE electrophoresis (8%, at 100 V for 90 min) followed by transfer to a PVDF membrane (BioRad). Membranes were washed in Tris-buffered saline (TBS) + Tween (TBST; 0.05% Tween 20), blocked for 1 h in TBST + 5% milk at room temperature, and incubated at 4°C overnight with α1H polyclonal (1:2,000), α1G monoclonal (1:500), TRPC1 monoclonal (1:1,000), or TRPC5 polyclonal (1:1,000) antibody. Incubation with goat anti-rabbit horseradish peroxidase (HRP)- or anti-mouse HRP-conjugated secondary antibodies was done at room temperature (1:2,000; Pierce). Blots were exposed to developing agent (Supersignal Femto Dura, Pierce) before analysis with a ChemiDoc MP System (BioRad).

### Electrophysiology

Whole-cell patch clamp recordings were performed from WT cultured hypothalamic neurons from 8-10 DIV. Square protocols to obtain I-V curves and T-type component subtraction were done as described in (Paula Patricia Perissinotti et al., 2015) and (Bean, 1985). Resting membrane potential (V_M_) and APs were recorded in external solution containing (in mM) 135 NaCl, 5 KCl, 2 CaCl_2_, 1 MgCl_2_, 10 HEPES, 10 glucose and intracellular solution containing (in mM) 110 K-gluconate, 20 KCl, 2 MgCl_2_, 1 EGTA, 10 HEPES, 2 ATP-Mg, 0.25 GTP-Li and 10 phosphocreatine-Tris. Pipette resistances were 3.2-4.5 MΩ. Cells with series resistance (*R_s_*) <20 MΩ were used; *R_s_* was compensated online (>80%). Data was acquired with and analyzed with pClamp 10 software (Molecular Devices). Cell capacitance was measured as described (Paula Patricia Perissinotti et al., 2015). Drugs: NNC-550396 dihydrochloride and Lep were purchased from Tocris (Bristol, UK).

#### Resting membrane potential

V_M_ was recorded in continuous trace mode without current injection for 20 s, and averaged; voltages were corrected for liquid junction potentials.

#### Input resistance

Whole cell input resistance (*R_in_*) was determined from V_M_= −60 mV in response to a −60 pA current step.

#### Membrane excitability

AP discharges were triggered by four consecutive depolarizing ramps at 20 and 40 pA/s rates from V_H_= −75 mV for 1.5 s. (Balasubramanyan, Stemkowski, Stebbing, & Smith, 2006; Lu et al., 2006). The rheobase was determined as the minimum amount of current required for firing an action potential from V_H_= −75 mV.

#### Single APs

APs were evoked from V_H_= −50 or −75 mV by a 3 ms-long rectangular pulse in steps of 50 pA.

#### Post inhibitory rebound (PIR) response

PIR protocols were well tolerated by most of the neurons tested in our culture. APs were evoked after 150 ms hyperpolarizing steps from −100 pA to 0 pA (Δ20 pA) from V_H_=-60 mV.

#### Steady state current

Normalized steady-state current (Istst) was calculated from the formula I= G* (V-E_Nernst(ion)_), where the channel conductance G is multiplied by the channel’s open probability and availability. Specifically, normalized steady-state current (Istst) was calculated with the steady stated formula Iss(V)= G*I/ImaxSSA(V)*I/ImaxSSI(V)*(V-40 mV), where V is the test voltage, G is the channel conductance (arbitrarily set at 1) and I/ImaxSSA and I/ImaxSSI are obtained experimentally from the LVA steady state curves adjusted to a Boltzman equation; 40 mV is the experimentally equilibrium potential for calcium.

### Statistical analysis

Results are presented as mean ± SEM. Statistical analysis was performed with the Sigma Plot 11 Software. Statistical significance (*P*< 0.05) was determined using Student’s t-test, unless otherwise noted.

## Abbreviations

AgRP: agouti-related peptide
AP: action potential
DIV: days *in vitro*
KLHL1: Kelch-like 1
Lep: Leptin
LRb: Leptin receptor
LVA: low-voltage activated
NPY: neuropeptide Y
PIR: post-inhibitory rebound
POMC: proopiomelanocortin positive neurons
TRPC: Transient Receptor Potential Cation
TRPC5: Transient Receptor Potential Cation 5

## Acknowledgements

We thank all members of the Piedras laboratory, Drs. O’Connell and Don Carlos for helpful input and comments. This paper is based upon work supported by the National Science Foundation under Grant no. 1022075 (EPR).

## Author contributions

E.S.P.R. conception of research; E.S.P.R., E.M.H and P.P.P. designed the experiments; P.P.P., E.M.H and E.S.P.R. performed experiments; E.S.P.R supervised the experiments; P.P.P., E.M.H and E.S.P.R analyzed data, interpreted results of experiments; P.P.P, E.M.H. and E.S.P.R prepared figures; E.S.P.R and P.P.P wrote the manuscript; P.P.P and E.S.P.R edited and revised the manuscript. E.S.P.R approved the final version of the manuscript.

## Conflict of Interest Statement

The authors declare that the research was conducted in the absence of any commercial or financial relationships that could be construed as a potential conflict of interest.

## Notes

### Competing Interest Statement

The authors have declared no competing interest.

